# Unveiling KRAS Mutant Structures with AlphaFold 3: New Avenues for Targeted Cancer Therapy

**DOI:** 10.1101/2025.05.16.654380

**Authors:** Guanxue Zhao, Yu Zhao

## Abstract

Despite recent advances in targeting the “undruggable” KRAS (Kirsten rat sarcoma viral oncogene homologue) protein, much of its structural information remains elusive. Utilizing a “branch-pruning” strategy, this study systematically mutated the Switch I and II regions of KRAS, predicting mutants’ structures with AlphaFold 3 and analyzing their structure differences with KRAS wildtype using PyMOL. The analysis of 494 mutants showed that over 64% had root-mean-square deviation (RMSD) scores below 0.200, indicating minor structural changes compared to wildtype KRAS. Notably, Switch II exhibited greater conformational variability, suggesting its potential role in oncogenic signaling. This work found that Q61 mutants showed limited structural changes, but nearby sites like E63 and M67 were more dynamic. Highly dynamic sites D38 (on Switch I) and E76/G60 (on Switch II) were identified, potentially affecting KRAS dimerization and post-translational modifications in oncogenesis. Mutants like D31I, R73C, and T74Q induced conformational changes, yet their biological functions are still unknown, possibly impacting effector binding and allosteric communication. These findings could reveal cryptic pockets for drug targeting, deepening our understanding of KRAS’s role in cancer.

## Introduction

### 1.1 KRAS structure and function

KRAS protein is a small GTPase and its mutations are implicated in 25-30% of all human cancers^1^. Among different cancer types, the top three most KRAS driven cancers include pancreatic cancers (73%), colorectal cancers (40%), and lung cancers (19%)^1^.

RAS family proteins, encoded by *KRAS (Kirsten rat sarcoma viral oncogene), HRAS (Harvey rat sarcoma viral oncogene)* and *NRAS (Neuroblastoma RAS viral (v-ras) oncogene)*, are small, membrane-bound guanine nucleotide-binding proteins, act as molecular switches by cycling between GTP-bound (active) and GDP-bound (inactive) conformations. RAS proteins have a crucial role in the regulation of cell proliferation, differentiation and survival by signaling through several important pathways^2^.

KRAS proteins are activated by a guanine nucleotide-exchange factor (GEF) and inactivated by a GTPase-activating protein (GAP). In the presence of mitogenic signals, KRAS proteins recruit GEF, catalyzing GTP loading, transiting the active sites from an open to a closed conformation and promoting subsequent interactions with various effector proteins^3^. The dynamic active sites of GTP-bound KRAS proteins with closed conformation are capable of binding to a few different effectors and regulators proteins to influence signal transduction, including Phosphatidylinositol 3-kinase (PI3K), Stress-activated protein kinase-interacting protein 1 (SIN1) of mechanistic Target of rapamycin complex 2 (mTORC2), Rapidly accelerated fibrosarcoma (RAF) kinases, etc^3^. However, the affinities and selectivity of the different RAS proteins for any given effector have not been systematically analyzed^3^.

Owing to a splice variant (distinct forms of mRNA and proteins generated from a single gene through alternative splicing, which can lead to different functions and properties), *KRAS* encodes for two proteins, KRAS4A and KRAS4B, which differ only in the final C terminal (hypervariable region). In this research, we primarily focus on KRAS4A (UniProt ID: P01116), which contains 189 amino acids. The RAS family proteins have been discovered over 40 years with well-characterized structures. For KRAS protein, there are four main regions border the nucleotide-binding pocket: the phosphate-binding loop (P-loop, residues 10-17), Switch I (residues 30-38), Switch II (residues 60-76) and the base-binding loops (residues 116-120 and 145-147)^2^. Switch I and II regions differ in conformation between the GDP and GTP states and dynamically interact with RAS-binding partners. G60 and T35 sites from the two Switch regions act as springs anchored at the γ-phosphate of GTP of RAS protein^2^.

### 1.2 KRAS mutations and their activation

*KRAS* mutations are by far the most common lesions found in cancer, including pancreatic, colorectal, and lung cancers. Similar to other RAS isoform mutations, mutations in KRAS can impact the intrinsic or GEF-mediated GDP/GTP exchange rate, and the intrinsic or GAP-mediated hydrolysis rate^4,5^. Among currently identified *KRAS* mutant alleles, the three common mutational sites of KRAS proteins are located on residue 12, 13 in the P-loop and 61 in switch II^1^. Mutation of KRAS proteins displays strong site preference, with ∼80% of KRAS mutations occurring at residue G12, ∼15% at G13, and ∼5% at Q61^12^.

Most KRAS mutations are deficient for intrinsic hydrolysis. However, KRAS G12C maintains much of its intrinsic hydrolysis activity (which means an activity that is not catalyzed by any other protein or chemical). KRAS G12R, G13D, Q61H, Q61L, and A146T mutations have increased intrinsic exchange relative to wildtype, but G12C and G12V do not display large changes^6^. These impacts on intrinsic GTPase activity can lead to biological differences between point mutations activating KRAS^1^.

These results suggested that KRAS G12C retains intrinsic GTPase activity and is sensitive to GAP-induced hydrolysis, two functions that are necessary to maintain a pool of GDP-bound KRAS G12C. This mechanism was employed to design covalent inhibitors that bind to GDP-bound KRAS through cysteine at residue 12, locking the active site in an inactive conformation. Sotorasib, the first covalent inhibitor for KRAS G12C, has been approved for the treatment of patients with advanced lung adenocarcinoma expressing this mutant and who have failed at least one systemic pretreatment^7^. Unfortunately, resistance to G12C inhibition develops rapidly through a variety of mechanisms, including amplification of the mutant allele, mutations in downstream members of other signaling pathways, acquisition of oncogenic fusions, and/or trans-differentiation to an alternative cellular state and more common *de novo* mutations in KRAS itself^8^.

Based on the biological validation of functional differences occurred in the past 5 years, the core biochemical properties of KRAS mutants on its 189 amino acid protein sequence could be classified into several classes: class 1 (hydrolysis, only KRAS G12C mutant was included), class 2 (exchange,15 KRAS mutants were included), class 3 (hybrid, 3 KRAS mutants were included, mainly located in switch II region) and class 4 (To be determined (TBD), most mutants were classified in this group)^3^.

Class 4 KRAS mutants are usually located on the surface of the protein, distal from active regions, or do not directly interact with the guanine nucleotide. Although the biochemical properties of KRAS proteins with class 4 mutants are not well characterized (are referred to as TBD) and are found more often in germ-line variants of KRAS, it is possible that these amino acid substitutions might influence a wide range of KRAS functions. These functions include affinity for effector and regulatory proteins, rates of nucleotide exchange, and the ability of KRAS to attain the catalytically competent conformation^3^. After analyzing many germ-line KRAS variants, *Schubbert et al*. demonstrated that despite reduced hydrolysis and enhanced nucleotide exchange, germ-line KRAS variants generally have reduced effector binding affinity, such as RAF, to prevent the full activation of downstream pathways ^9^. Although germ-line mutations are not more frequently observed in cancer, exploring how these variants influence the activation of KRAS will certainly lead to a more holistic understanding of KRAS function and the relevance of drug development strategies that target these interactions.

### 1.3 Conformational dynamics of KRAS proteins

Recent progress in structural biology revealed that proteins’ structures are flexible and mobile, with parts of proteins moving on different timescales and to different degrees^2^. The creation of potent RAS inhibitors has been challenging due to the mechanism by which RAS engages with its downstream effector through direct protein-protein interactions, and the overactivation of RAS in cancer is linked to the dysfunction of its enzymatic activity. Consequently, for a RAS inhibitor to be effective, one might exploit some strategies, such as reducing the amount of RAS in the GTP-bound state, interfering with the RAS-GTP-effector complex formation, promoting the formation of inactive protein complexes, or lowering the concentration of RAS at the cell membrane^2^.

Hence, understanding protein architecture is vital for the drug discovery process, as the sites where small molecules bind can be dynamic in form, or a binding site may only become apparent upon interaction with small molecules, which was not noticeable in the unbound structure. Structural studies of RAS in the GDP state indicate increased flexibility in the switch regions, especially Switch II, relative to the GTP state^2^. *Gorfe et al*. concluded that although greater mobility in KRAS structures was identified in both GDP-bound mutants (such as KRAS G12C and KRAS Q61L) and GTP-bound wildtype, the GTP-bound KRAS structures are much more rigid in the switch II region^10^. Switch I/ II regions, a small, shallow, and polar pocket as the most important areas for interactions with GEFs, GAPs, and downstream effectors, are considered the most important drugging pocket on KRAS^17^.

The development of computational techniques helps scientists unravel the molecular mechanisms underlying KRAS mutants. By sequence alignment and clustering analysis between KRAS wildtype and its mutated variants (such as G12C, G12D, G12V, and G13D), higher conformational changes were identified in the α-helix in mutated variants of KRAS^11^. The RMSD score of KRAS wildtype ranges between 0.082-0.280, with an average RMSD of 0.182, suggesting a relative stability of the protein^11^.

Molecular dynamics simulations analysis of wildtype and KRAS Mutants revealed that the RMSD score range of KRAS mutants is within the wildtype range, indicating that the KRAS G12C, G12D, G12V, and G13D mutants do not significantly disrupt the stability^11^. However, this work only focused on known KRAS mutants and did not tap into a wider range of non-cancer-related KRAS sequences, especially those “TBD” mutants.

To understand how different RAS mutations would affect its binding with downstream effectors, *Phillipp et al*. developed a “branch-pruning” strategy (I36 and E37 sites on the RAS interface were substituted by other 19 amino acids in a sequential order) to generate different RAS mutations and employed AlphaFold 2 to predict the RAS-effector interactions^13^. This single amino□acid substitution strategy is also named “edgetic” (edge-specific genetic perturbations) to provide alternative molecular explanations for protein dysfunction in addition to gene loss^14^.

Despite its well-established role in oncogenesis, drug design targeting KRAS has historically been challenging due to its nearly spherical structure with no obvious binding site on the protein surface^16^. With the help of AlphaFold, we can have a comprehensive understanding of the impact of mutations on the structural dynamics and stability changes of GDP-bound KRAS proteins. It is also reported that harmless passenger mutations accumulated by cancer cells may impact proteome stability and lead to proteotoxic stress^15^. Motivated by *Phillipp et al*.’s work, here we use AlphaFold 3 to simulate all variants in KRAS Switch I/ II regions to advance our understanding of the structural biology of KRAS.

## Results

### Sequence alignment of KRAS wildtype protein structures from PDB and AlphaFold□3 prediction

AlphaFold□3 (AF3), the latest version of AlphaFold, was considered a powerful tool to predict protein structure with high accuracy for nearly all molecular types presented in the Protein Data Bank (PDB)^18^. We first aligned the protein structure from PDB (KRAS wildtype: UniProt ID: P01116) with the AF3’s prediction of KRAS wildtype structure (Fig. 1). The RMSD score was 0.177, indicating high accuracy of AF3 protein structure prediction.

**Fig. 1.**
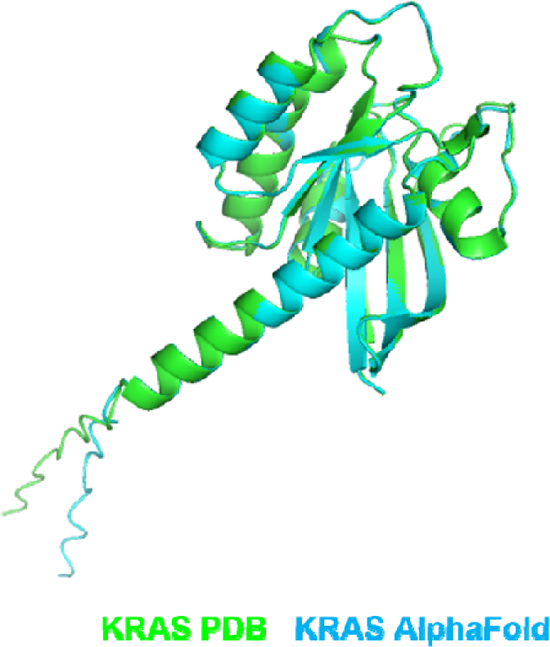
Structure alignment of KRAS wildtype obtained from PDB and AlphaFold 3 prediction (RMSD: 0.177)

A major constraint of AF3 models is their tendency to forecast rigid structures akin to those in the PDB, rather than the dynamic nature of biomolecular systems in a liquid environment, which impairs the precision of the modeled conformations^18^. *Abramson et al*. identified stereochemical inaccuracies, false positives, insufficient dynamics, and target-specific inaccuracies as the principal drawbacks of AF3 model reliability^18^. Although this RMSD score indicates improved areas for AF3 protein structure prediction, we believe that AF3 still satisfies our research goal of exploring structural variants on KRAS in this work.

### Sequence alignment of KRAS wildtype and mutated variants

To further study the impact of specific mutations on the protein’s structural stability, we focused on Switch I (site 30-38) and Switch II (site 60-76) and substituted each site with other 19 amino acids. After employing the systematic mutation strategy, we derived 494 KRAS mutants, and then aligned them with KRAS wildtype. The RMSD score of the alignment of KRAS wildtype and mutants varied between 0.163 (I36Y) -0.229 (G75L) and summarized in Table 1, with 39 mutants’ RMSD scores higher than 0.210. Fig. 2 shows a heatmap of all these 494 mutants based on different RMSD scores. The variations in RMSD detected between the wildtype KRAS and its mutated forms imply potential implications for the functional kinetics of these KRAS variants, offering crucial understanding of how particular mutations affect the protein’s structural robustness.

**Table 1.**
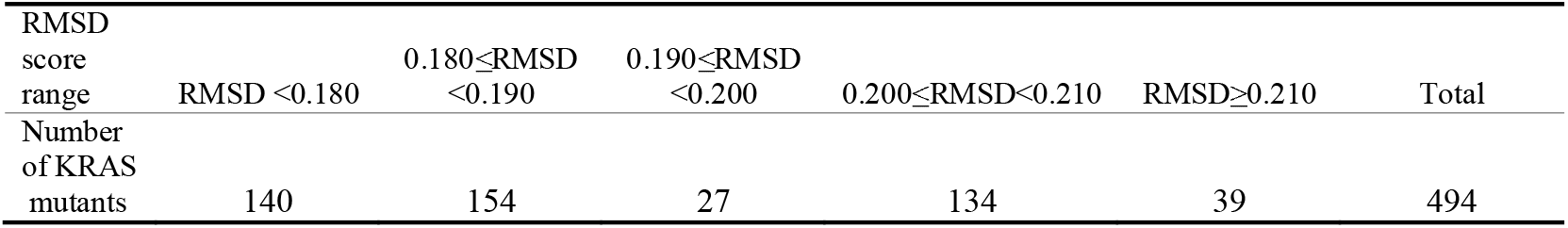
Distribution of KRAS mutants in different RMSD score range.

**Fig. 2.**
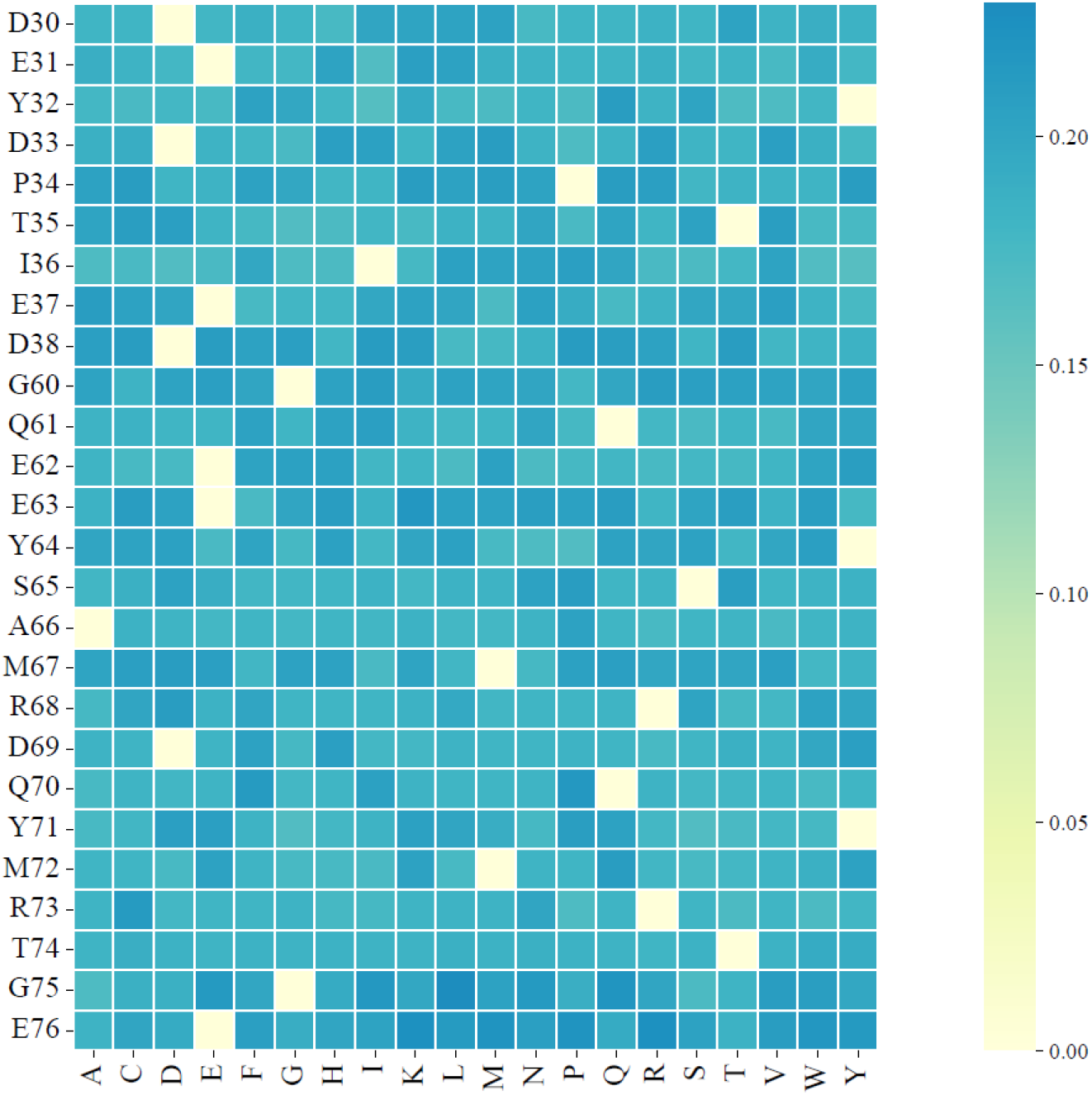
Heat map clustering of systemic mutations on KRAS Switch I (site 30-38) and Switch II (site 60-76) regions.

We also calculated the total RMSD score for each site on KRAS Switch I and II region, and highlighted D38 (on Switch I) and E76 (on Switch II) sties, displaying the highest total RMSD score of 3.744/ 3.915, respectively (Table 2). The Switch II region appeared to harbor a greater number of dynamic sites compared to the Switch I region, as evidenced by the identification of multiple sites with higher total RMSD scores than D38, including G60, G75, and E63. The elevated total RMSD scores suggest that these sites likely play a significant role in shaping the overall structural dynamics of the protein.

**Table 2.**
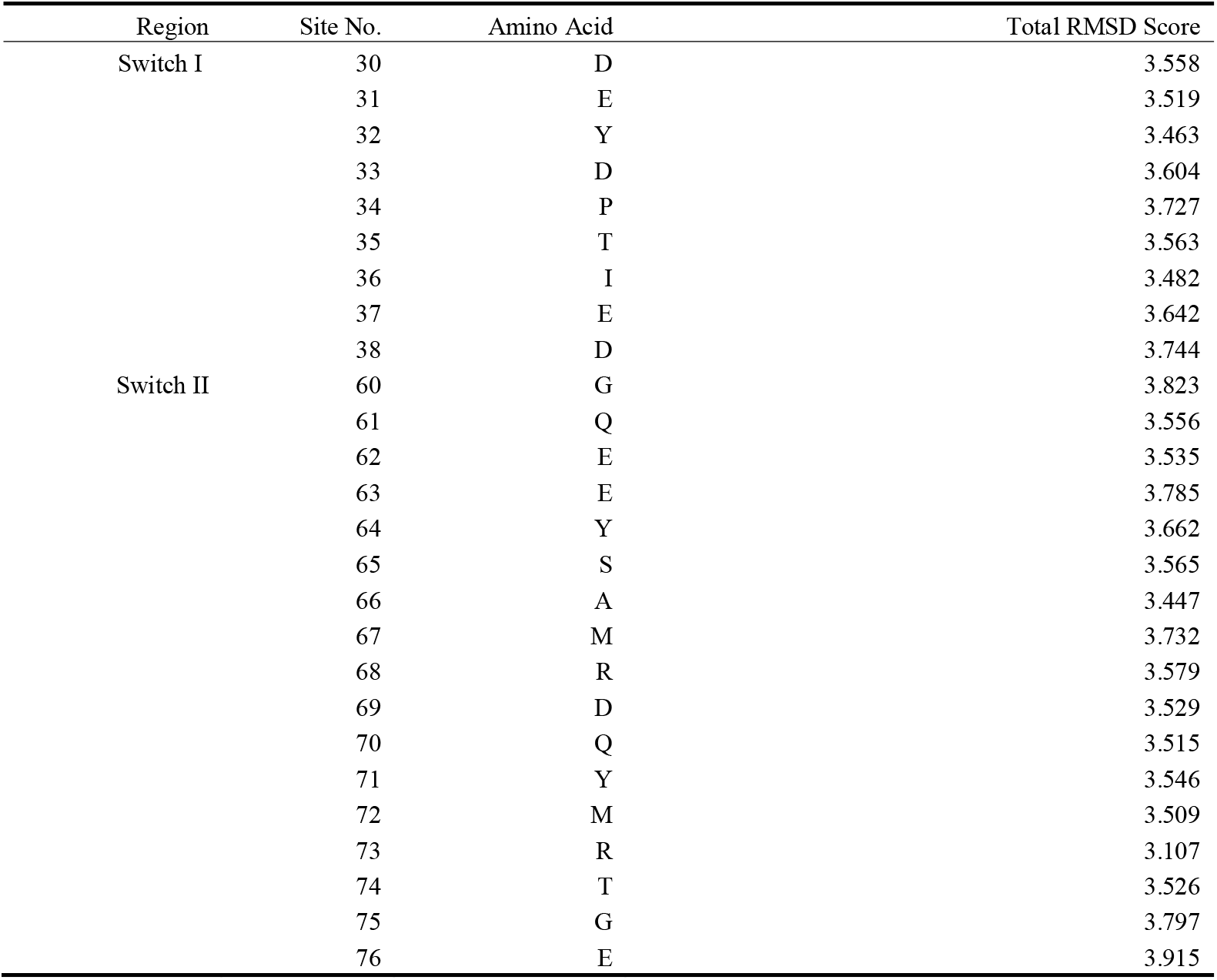
Summary of the total RMSD scores for KRAS mutations resulting from systematic amino acid substitutions in Switch I/ II regions.

### Visualized analysis of the alignment of KRAS wildtype and mutated variants

PyMOL provides a visualized comparison of the impacts of site mutations on KRAS conformation changes. We summarized a few typical visualized comparisons in Fig.3. RMSD score analysis of those 494 alignments suggests most mutations do not introduce significant changes in the protein structure compared to the wildtype, even the RMSD scores are higher than 0.210 (Fig. 3a). However, we still can identify relative conformation changes on some of the mutants, such as D31I mutant (RMSD=0.169), R73C mutant (RMSD=0.213) and T74Q mutant (RMSD=0.183) (Fig. 3b). Given that mutations at position E76 induced more pronounced conformational dynamics, we have depicted several of these mutants in Fig. 3c, namely the E76C, E76S, and E76T variants.

**Fig. 3.**
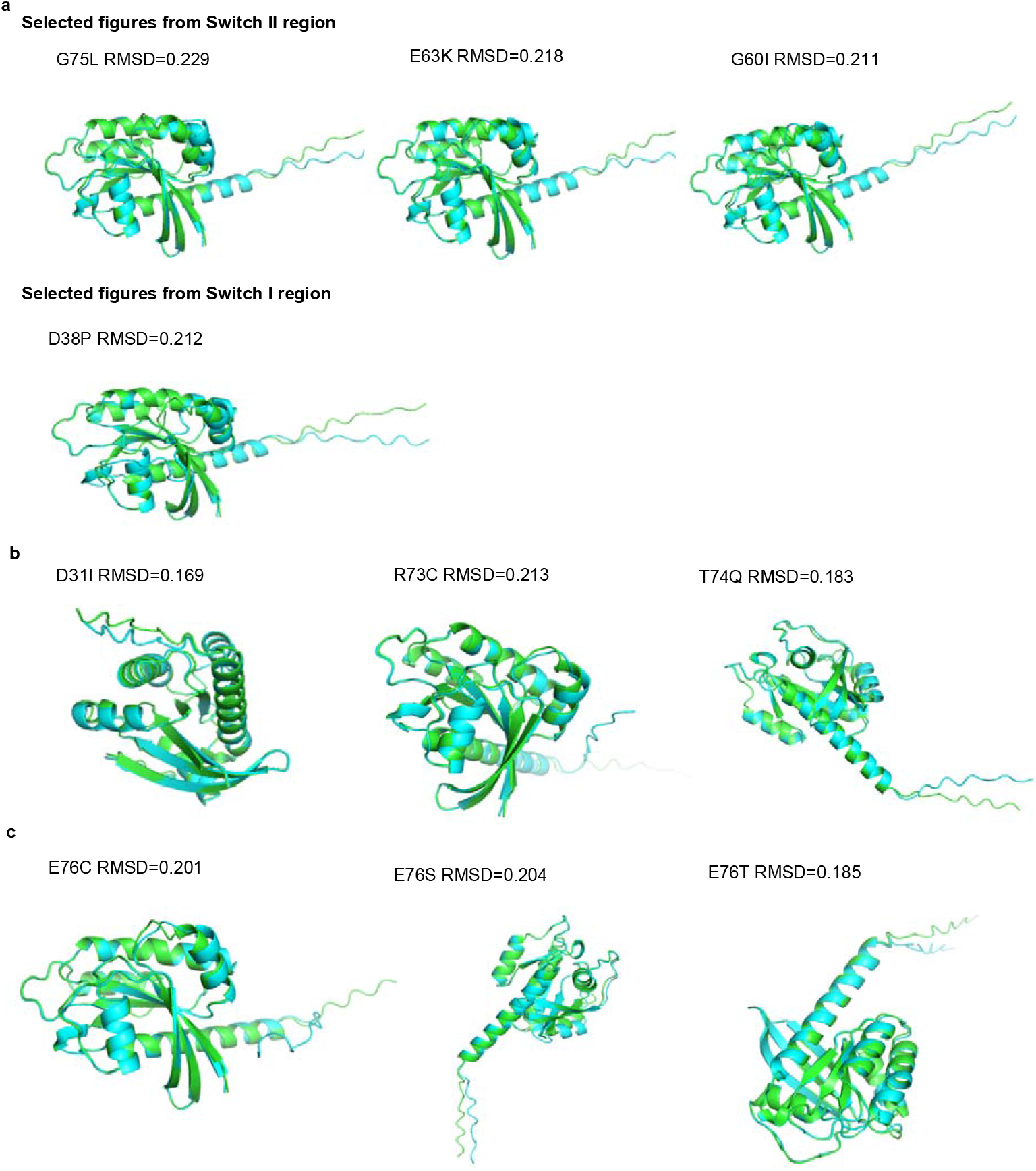
Structure alignment of KRAS mutants (Blue) with KRAS wild type (Green). a) Selected structure alignment figures from sites with higher total RMSD Score, such as D38 (Switch I) and G60/ E63/ G75 (Switch II). b) Structure alignment figures showed different patterns compared to those displayed in Fig.3a. c) Pronounced conformational dynamics identified from E76C, E76S, and E76T mutants.

## Discussion

KRAS has a nearly spherical structure with no obvious binding site, making it historically difficult to target small molecules. Advances in computational techniques, such as AF3, have enabled deeper insights into KRAS structural variations and potential druggable sites. In this work, we created systematic single-site mutations primarily focused on the dynamic regions of KRAS, such as Switch I (residues 30-38) and Switch II (residues 60-76). KRAS mutant structures predicted by AF3, and aligned with wild-type KRAS using PyMOL to calculate RMSD. Our results show that over 64% of the total 494 mutants’ RMSD scores fall below 0.200 and the highest score is only at 0.229, indicating moderate conformational fluctuations triggered by most signal amino acid substitutions. These results are not surprising, as even the mutation on the most wildly studied KRAS residue 12 (G12C, G12V, G12D, and G13D) which has been proven to have a crucial role in cancer development didn’t dramatically affect protein structure stability compared to the wildtype^11^.

Interestingly, sites on the Switch II region displayed a higher total RMSD Score than those on the Switch I region, suggesting more dynamics in mutant structure changes on the Switch II region. Switch II region contributed to the rigid binding of KRAS with GTP and was more dynamic in structural changes compared to other functional regions of KRAS^10^. Nevertheless, Q61, one of the three most frequently occurring oncogenic mutation sites, did not register a high total RMSD score, with a value of only 3.556. Surprisingly, our study revealed notably high total RMSD scores for E63 (3.785) and M67 (3.732). This result can be accounted for by *Vatansever et al*.’s observations concerning an H-bond network within the Switch II region: 1) R68 engages in bonds with E63, S65, Y71, and M72; 2) M67 is connected to Q70; and 3) Q61 is linked to E63, suggesting that mutations at these sites may alter the local conformation and dynamics of KRAS^23^. *Vatansever et al*. concluded that through GTP hydrolysis impairment, the activation of GTP-bounded KRAS G12D mutant signaling was probably due to the disruption of the H-bond network within Switch II coupled with the deviation of central Switch II region toward α3 helix^23^. Furthermore, our findings indicate that the Q61 mutant undergoes a consistent conformational shift in KRAS, aligning with prior reports on the oncogenic mechanisms of the KRAS Q61 mutant through a displacement of α3(Y96) away from Switch II (G60) during GTP-to-GDP transition^24^.

We further expand on the biological implications of total RMSD results for KRAS wildtype and mutated variants, especially for D38 and E76/ G60. As D38 is in the β interface and is exclusively involved in the α-β interaction^20^, *Lee et al*. concluded that single-point mutations at either the α-α or α-β dimer interface are sufficient to reduce dimerization as proved by the D38K or E168R mutant model in their study^19^. Besides the mutation mechanism of impairing GTP hydrolysis and locking KRAS protein in an active GTP-bound state, affecting KRAS dimerization has recently emerged as an additional layer of KRAS mutant-mediated oncogenesis.

Although not recognized as a popular cancer-driven mutation, G60 on KRAS was reported as one of the germline KRAS mutations, affecting phenotype development of diseases, such as Noonan syndrome (NS), cardio-facio-cutaneous syndrome (CFCS), and Costello syndrome (CS). The mutation of KRAS G60 to arginine has been reported to cause the impairment of almost all biochemical and functional properties, including nucleotide binding, GTP hydrolysis, and interaction with effectors^21^. *Lothar et al*. have elucidated that the pivotal role of G60 in mediating the critical conformational transitions between the active and inactive states of germline KRAS mutants could be explained by its strategic structural position, as G60 substitutions can directly influence nucleotide binding and hydrolysis in the GTP-bound form, and this residue, which interacts with the CDC25 domain of SOS1, becomes solvent-exposed upon binding GDP^22^.

In our study, E76 emerged as the most dynamically active site within both Switch I and II regions, intriguingly positioned at the terminus of Switch II. We highlight that E76 has hitherto been largely overlooked in research, which may be due to the presumption that this area could play a role in modulating KRAS-GAP interactions. E76 mutations were found to cause cell transformation to a tumor phenotype *in vitro* and *in vivo*, and synergize with Ras G12 mutations to promote tumor growth^27^. Examining the alignment images of the E76 mutants, we underscored the distinct conformational shifts observed in the E76C, E76S, and E76T variants, which set them apart from other E76 mutants, in the absence of any documented biological implications.

While no documented biological significance was found for the conformational alterations induced by the D31I, R73C, and T74Q mutants, we hypothesize that these mutations may conceal critical, yet unexplored functions related to effector binding or influencing the allosteric communication with other residues. The functionality of KRAS is finely tuned by an array of post-translational modifications, including farnesylation, palmitoylation, and phosphorylation, among others. Several distal sites within KRAS have been implicated in modulating GEF-mediated nucleotide exchange, in concert with specific residues on the Switch I/ II regions, such as K104, which interacts with M72, R73, and G75 on Switch II, influencing KRAS protein acetylation. We propose that certain mutants within the Switch I/ II regions could disrupt these interaction networks and modifications, potentially explaining their previously overlooked importance in research. The structural alterations of the protein resulting from various mutations will aid in uncovering the hidden pockets within the KRAS protein structure, as cryptic pockets are likely less conserved compared to orthosteric sites^25^. Leveraging PocketMiner, *Arthur et al*. reported to identify cryptic pockets for over half of the proteins which were previously be overlooked as drug targets^26^.

In conclusion, this research has established a database cataloging sequence-based structural variations of KRAS in the Switch I and II regions through computational methods, visually documenting the impact of KRAS mutants. Utilizing the robust AF3 platform, our study offers significant insights that are poised to enhance the treatment of KRAS mutated cancers. Within this context, we conducted preliminary analyses of the structural features of various KRAS mutants using RMSD scores. The results identify distinct conformational dynamics across the mutated KRAS proteins, particularly at the D38 and E76/G60 sites, shedding light on their potential functional implications.

Moving forward, the scope of this research can be broadened to examine additional regions of the KRAS sequence, aiming to uncover further conformational alterations brought about by KRAS mutations. The strategy of sequential site mutations can also be employed to elucidate the structural dynamics of known KRAS mutations, particularly when considering various amino acid substitutions, such as G12 mutations in conjunction with other mutations along the KRAS sequence. Moreover, partnering with experimental methodologies will be instrumental in validating the robustness of our findings, as we continue to advance our investigative efforts in this field.

## Methods

### Protein alignment and Visualization

All software applications were installed and operated on a personal computer with a Windows 10 operating system, equipped with an Intel i5-1135G7 processor and 8 CPU cores.

The protein sequence data of KRAS specific to humans was acquired from the AlphaFold^29^ database by searching for the UniProt ID: P01116.

KRAS mutant structure was predicted by AlphaFold 3: https://alphafoldserver.com/.

The results from AlphaFold 3 were visualized and aligned with wildtype KRAS using PyMOL, version 2.5.7 for Windows-x86_64.

### KRAS Protein mutation strategy

Employing a “branch-pruning” approach^13^, we have developed models for single-site KRAS mutants, focusing on the functional architecture of KRAS, namly the Switch I (residue 30-38) and Switch II (residue 60-76) regions, encompassing a total of 26 mutation sites. We replaced each site with 19 other amino acids and derived 494 KRAS mutants, whose protein structure were predicted by AlphaFold 3. KRAS mutants’ structures were loaded into PyMOL and aligned with KRAS wildtype.

## References

[1] Ash, L.J.; Busia-Bourdain, O.; Okpattah, D.; Kamel, A.; Liberchuk, A.; Wolfe, A.L. KRAS: biology, inhibition, and mechanisms of inhibitor resistance. Curr. Oncol. 2024, 31, 2024–2046.

[2] Ostrem, J.; Shokat, K. Direct small-molecule inhibitors of KRAS: from structural insights to mechanism-based design. Nat Rev Drug Discov 2016, 15, 771–785.

[3] Johnson C.; Burkhart D. L.; Haigis K. M. Classification of KRAS-activating mutations and the implications for therapeutic intervention[J]. Cancer discovery 2022, 12(4), 913–923.

[4] Forbes, S. A.; Bindal, N.; Bamford, S.; Cole, C.; Kok, C. Y.; Beare, D.; Futreal, P. A. COSMIC: mining complete cancer genomes in the catalogue of somatic mutations in cancer. Nucleic acids research 2010, 39(Suppl_1), D945–D950.

[5] Prior, I. A.; Lewis, P. D.; Mattos, C. A. Comprehensive survey of Ras mutations in cancer. Cancer Res. 2012, 72, 2457–2467.

[6] Moore, A.R.; Rosenberg, S.C.; McCormick, F.; Malek, S. RAS-targeted therapies: is the undruggable drugged? Nat. Rev. Drug Discov. 2020, 19, 533–552.

[7] Skoulidis, F.; Li, B. T.; Dy, G. K.; Price, T. J.; Falchook, G. S.; Wolf, J.; Govindan, R. Sotorasib for lung cancers with KRAS p. G12C mutation. New England Journal of Medicine 2021, 384(25), 2371–2381.

[8] Misale, S.; Di Nicolantonio, F.; Sartore-Bianchi, A.; Siena, S.; Bardelli, A. Resistance to anti-EGFR therapy in colorectal cancer: from heterogeneity to convergent evolution. Cancer Discov 2014, 4, 1269–80.

[9] Schubbert, S.; Bollag, G.; Lyubynska, N.; Nguyen, H.; Kratz, C. P.; Zenker, M.; Shannon, K. Biochemical and functional characterization of germ line KRAS mutations. Mol Cell Biol 2007, 27, 7765–70.

[10] Gorfe, A. A.; Grant, B. J.; McCammon, J. A. Mapping the nucleotide and isoform-dependent structural and dynamical features of Ras proteins. Structure 2008, 16, 885–896.

[11] Mir, S. A.; Nayak, B.; Aljarba, N. H.; Kumarasamy, V.; Subramaniyan, V.; Dhara, B. Exploring KRAS protein dynamics: an integrated molecular dynamics analysis of KRAS wild and mutant variants. ACS Omega 2024, 9 (28), 30665–30674.

[12] Silverman, I.; Gerber, M.; Shaykevich, A.; Stein, Y.; Siegman, A.; Goel, S.; Maitra, R. Structural modifications and kinetic effects of KRAS interactions with HRAS and NRAS: an in silico comparative analysis of KRAS mutants. Front. Mol. Biosci. 2024, 11, 1436976.

[13] Philipp, J.; Khrisina, K. Structure-based prediction of Ras-effector binding affinities and design of “branchegetic” interface mutations. Structure 2023, 31(7), 870–883.

[14] Zhong, Q.; Simonis, N.; Li, Q.R.; Charloteaux, B.; Heuze, F.; Klitgord, N.; Tam, S.; Yu, H.; Venkatesan, K.; Mou, D. Edgetic perturbation models of human inherited disorders. Mol. Syst. Biol. 2009, 5, 321.

[15] Nagel, R.; Semenova, E. A.; Berns, A. Drugging the addict: non-oncogene addiction as a target for cancer therapy. EMBO Rep. 2016, 17(11), 1516–31.

[16] Pantsar, T. The current understanding of KRAS protein structure and dynamics. Comput Struct Biotechnol J. 2020, 18, 189–98.

[17] Kessler, D.; Gmachl, M.; Mantoulidis, A.; Martin, L. J.; Zoephel, A.; Mayer, M.; McConnell, D. B. Drugging an undruggable pocket on KRAS. Proceedings of the National Academy of Sciences 2019, 116(32), 15823–15829.

[18] Abramson, J.; Adler, J.; Dunger, J.; Evans, R.; Green, T.; Pritzel, A.; Jumper, J. M. Accurate structure prediction of biomolecular interactions with AlphaFold 3. Nature 2024, 1–3.

[19] Lee, K. Y.; Enomoto, M.; Gebregiworgis, T.; Gasmi-Seabrook, G. M.; Ikura, M.; Marshall, C. B. Oncogenic KRAS G12D mutation promotes dimerization through a second, phosphatidylserine– dependent interface: a model for KRAS oligomerization. Chemical Science 2021, 12(38), 12827–12837.

[20] Nan, X.; Tamguney, T. M.; Collisson, E. A.; Lin, L. J.; Pitt, C.; Galeas, J.; Lewis, S.; Gray, J. W.; McCormick, F.; Chu, S. Ras-GTP dimers activate the mitogen-activated protein kinase (MAPK) pathway. Proc. Natl. Acad. Sci. U. S. A. 2015, 112, 7996–8001.

[21] Cirstea, I.C.; Kutsche, K.; Dvorsky, R.; Gremer, L.; Carta, C.; Horn, D.; Roberts, A. E.;Lepri, F.; Merbitz-Zahradnik, T.; Konig, R.; Kratz, C. P.; Pantaleoni, F.; Dentici, M. L.; Joshi, V. A.; Kucherlapati, R. S.; Mazzanti, L.; Mundlos, S.; Patton, M. A.; Silengo, M. C.; Rossi, C.; Zampino, G.; Digilio, C.; Stuppia, L.; Seemanova, E.; Pennacchio, L. A.; Gelb, B. D.; Dallapiccola, B.; Wittinghofer, A.; Ahmadian, M. R.; Tartaglia, M.; Zenker, M. A restricted spectrum of NRAS mutations causes noonan syndrome. Nat Genet 2010, 42, 27–29.

[22] Gremer, L.; Merbitz-Zahradnik, T.; Dvorsky, R.; Cirstea, I.C.; Kratz, C.P.; Zenker, M.; Wittinghofer, A.; Ahmadian, M.R. Germline KRAS mutations cause aberrant biochemical and physical properties leading to developmental disorders†. Hum. Mutat. 2011, 32, 33–43.

[23] Vatansever, S.; Erman, B.; Gümüş, Z. H. Oncogenic G12D mutation alters local conformations and dynamics of K-Ras. Sci Rep 2019, 9, 11730.

[24] Prakash, P.; Sayyed-Ahmad, A.; Gorfe, A. A. The role of conserved waters in conformational transitions of Q61H K-ras. PLoS Comput Biol 2012, 8, e1002394.

[25] Ivetac, A.; Andrew McCammon, J. Mapping the druggable allosteric space of g-protein coupled receptors: a fragment-based molecular dynamics approach. Chem. Biol. Drug Des. 2010, 76, 201–217.

[26] Meller, A.; Ward, M.; Borowsky, J. Predicting locations of cryptic pockets from single protein structures using the PocketMiner graph neural network. Nat Commun 2023, 14, 1177.

[27] Guo, Z.; Oakes, S.; Lu, Y.; Sugita, M.; Franklin, W.; Franzusoff, A. Oncogenic synergy and a newly identified transforming Ras mutation in human cancer. Cancer Research 2007, 67(9_Supplement), LB-345.

[28] Yang, M. H.; Nickerson, S.; Kim, E. T.; Liot, C.; Laurent, G.; Spang, R.; Haigis, K. M. Regulation of RAS oncogenicity by acetylation. Proceedings of the National Academy of Sciences of the United States of America 2012, 109, 10843–8.

[29] Jumper, J.; Evans, R.; Pritzel, A.; Green, T.; Figurnov, M.; Ronneberger, O.; Tunyasuvunakool, K.; Bates, R.; Žídek, A.; Potapenko, A.; Bridgland, A.; Meyer, C.; Kohl, S. A. A.; Ballard, A. J.; Cowie, A.; Romera-Paredes, B.; Nikolov, S.; Jain, R.; Adler, J.; Back, T.; Petersen, S.; Reiman, D.; Clancy, E.; Zielinski, M.; Steinegger, M.; Pacholska, M.; Berghammer, T.; Bodenstein, S.; Silver, D.; Vinyals, O.; Senior, A. W.; Kavukcuoglu, K.; Kohli, P.; Hassabis, D. Highly accurate protein structure prediction with AlphaFold. Nature 2021, 596 (7873), 583–589.

